# RNA sequence and structure determinants of Pol III transcriptional termination in human cells

**DOI:** 10.1101/2020.09.11.294140

**Authors:** Matthew S. Verosloff, William K. Corcoran, Taylor B. Dolberg, Joshua N. Leonard, Julius B. Lucks

## Abstract

The precise mechanism of transcription termination of the eukaryotic RNA polymerase III (Pol III) has been a subject of considerable debate. Although previous studies have clearly shown that at the end of RNA transcripts, tracts comprised of multiple uracils are required for Pol III termination, whether upstream RNA secondary structure in the nascent transcript is necessary for robust transcriptional termination is still subject to debate. We sought to address this directly through the development of an *in cellulo* Pol III transcription termination assay using a synthetic biology approach. Specifically, we utilized the recently developed Tornado expression system and a stabilized Corn RNA aptamer to create a Pol III-transcribed RNA that produces a detectable fluorescent signal when transcribed in human cells. To study the effects of RNA sequence and structure on Pol III termination, we systematically varied the sequence context upstream of the aptamer and identified sequence characteristics that enhance or diminish termination. We found that in the absence of predicted secondary structure, only poly-U tracts longer than then the average length found in the human genome (4–5 nucleotides), efficiently terminate Pol III transcription. We found that shorter poly-U tracts could induce termination when placed in proximity to secondary structural elements, while secondary structure by itself was not sufficient to induce termination. These findings demonstrate a key role for sequence and structural elements within Pol III-transcribed nascent RNA for efficient transcription termination, and demonstrate a generalizable assay for characterizing Pol III transcription in human cells.

## Introduction

Cells require mechanisms for precisely terminating transcribed RNAs at the appropriate genetic loci in order to maintain proper genetic regulation and minimize undesired expression of downstream genomic regions [1-3]. The process of transcription termination in prokaryotes is well understood [4], but there remain questions regarding the mechanisms of termination in eukaryotes [1, 5-8]. This knowledge gap is particularly substantial for transcription mediated by eukaryotic RNA polymerase III (Pol III), which transcribes non-coding RNAs such as the 5S ribosomal RNA, tRNAs, snRNA, and a variety of miRNA [9, 10]. Given the important roles played by these classes of RNA in health and disease processes [11, 12], elucidating the mechanisms of transcription termination could provide insights into these import components of cellular regulation.

Pol III transcriptional termination occurs when the transcribing polymerase reaches a stretch of adenosines which is encoded into the nascent RNA as a poly-uracil (poly-U) tract [4]. The average lengths of these genomic tracts vary across eukaryotic species, with an average of 5–7 uracil nucleotides (nt) within the genome of *Schizosaccharomyces pombe* (*S. pombe*), 6–9 nt in *Saccharomyces cerevisiae* (*S. cerevisiae*) and 4–5 nt in humans [13]. In this regard, eukaryotic Pol III termination signals are similar to those employed in the bacterial intrinsic termination mechanism, which also occurs at a poly-U stretch [14]. In bacterial transcriptional termination, weak interactions between the A-U bases within the nascent RNA-DNA template hybrid signal the elongating RNA polymerase (RNAP) to transition into a pause conformation, and the resulting structural rearrangement contributes to the dissociation of the transcription complex from the template.

In addition to the poly-U tract, RNA secondary structure has been shown to play a role in transcription termination mechanisms [14]. For example, bacterial intrinsic transcriptional terminators require RNA secondary structures immediately adjacent to the poly-U tract, with more stable secondary structures contributing to greater termination efficiency [15, 16]. However, the necessity of RNA secondary structure for eukaryotic Pol III termination is currently debated [6, 7, 17-20]. Some reports indicate that Pol III efficiently terminates only when RNA structural elements are proximal to a poly-U tract [6, 17, 20], while other reports indicate that secondary structure is dispensable and Pol III termination is not enhanced in its presence [18, 21]. Most studies have utilized *in vitro* transcription assays using reassembled purified components of the *S. cerevisiae* Pol III transcription complex on DNA templates designed to contain various RNA sequence and structure contexts. In these assays, transcription components were first assembled into the full complex by loading on DNA templates, and then tested for their ability to read through poly-U tracts in the presence or absence of upstream RNA secondary structure [22]. The resulting transcript lengths were read out using radiolabeling and gel electrophoresis to determine whether or not upstream sequence and structural contexts were sufficient to terminate transcription at loci of interest. Even while using similar experimental procedures, several different groups observed different results as to whether RNA secondary structural elements adjacent to the poly-U tract enhances termination efficiency [6, 7, 17, 18]. It is not yet clear why these reports reached seemingly disparate conclusions. Possible explanations include subtle differences in enzyme preparations leading to the presence or absence of currently unknown termination determinants [18]. It is also possible that mechanisms of transcription termination significantly varies across eukaryotic species, and there indeed exists species-specific variations within the Pol III enzyme’s subunits that are known to influence transcription termination [21, 23]. To provide additional insight into this important question, we decided to address a gap in these observations by systematically studying the role of RNA secondary structure in transcription termination in human cell lines.

Here we present an assay for interrogating RNA sequence and structure determinants required for efficient Pol III termination in human cells. Specifically, we adapted an *in cellulo* Pol III transcriptional reporter system based on fluorescent RNA aptamers that are active in human cell lines [24]. RNA aptamers are structural motifs capable of binding to target ligands with high specificity [25]. Here we utilized the Tornado system which contains the Corn aptamer embedded within RNA transcripts for direct reporting of Pol III transcriptional activity (**Fig. 1A**). Placing specific RNA sequences upstream of the Tornado reporter system allowed us to assess the impact of these sequences on transcription efficiency. This system enabled us to determine how various transcript characteristics, including poly-U tract length and the presence and location of predicted RNA structural elements, influence Pol III termination efficiency in human cells. We anticipate that this work will further clarify the mechanism of Pol III transcription termination and enable the forward design of synthetic variants for precise control of Pol III expression in human cells.

**Figure 1:**
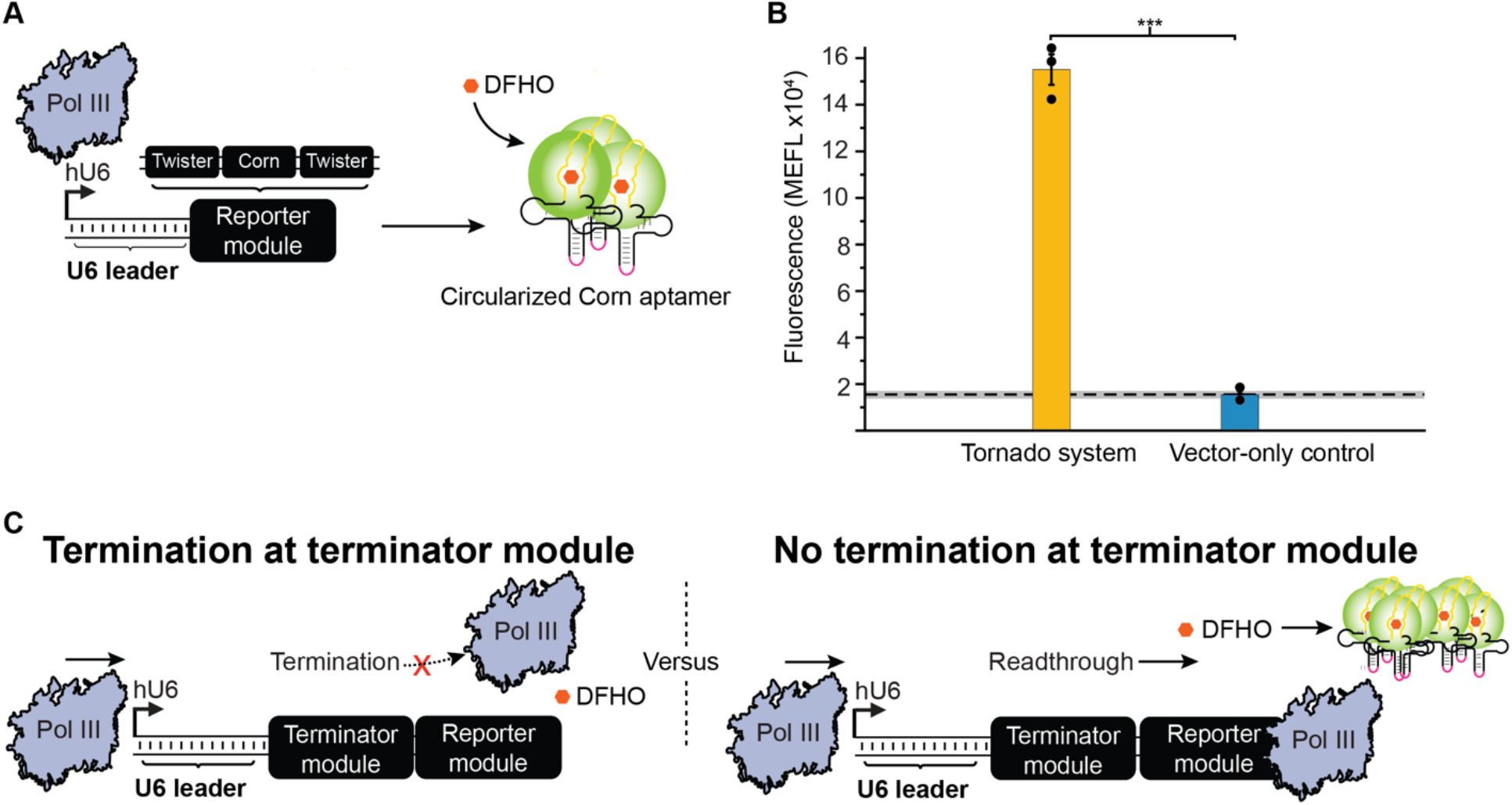
Development of an assay for measuring Pol III termination in human cells. **A)** Schematic overview of the construct design of the Tornado system [26]. A human U6 promoter drives Pol III transcription of a ‘Reporter module’, comprising a Corn aptamer (yellow line) which is embedded within a tRNA scaffold and flanked by two twister self-cleaving ribozymes. The ribozymes’ self-cleavage results in a 5’ hydroxyl and a 3’ end consisting of a 2’,3’-cyclic phosphate which is recognized and ligated (pink line) by the endogenous protein RtcB, increasing its stability. Ribozyme self-cleavage and ligation leaves a circularized Corn aptamer that is stabilized by the tRNA scaffold. When DFHO is bound to the Corn aptamer, this dye becomes fluorescent. **B)** Tornado enables quantification of Pol III transcription *in cellulo*. Colored bars represent the average of 3 biological replicates (circles). The dashed line indicates the average signal from the vector-only (negative) control cells, while the grey horizontal bar represents the standard error of the mean (S.E.M) of the signal from these cells; this convention is applied in subsequent figures. Statistical significance was measured using a one-tailed heteroscedastic Welch’s t-test (***p < 0.001). Error bars represent the S.E.M. **C)** Termination modules were introduced in this study to investigate the effect of different RNA sequences and structures on Pol III termination. Shown here are expected assay outcomes as a function of termination (left) or readthrough (right) at the Terminator module.

### An Assay for Quantifying Pol III Transcription Termination in Human Cells

To investigate the Pol III termination mechanism *in cellulo*, we first sought to develop a method that could quantitatively characterize the abundance of Pol III-generated transcripts within a human cell line. We started with previous work which used the fluorescent RNA aptamer, Corn, to study the subcellular localization of Pol III transcripts [24]. When transcribed, Corn forms a secondary structure that binds the ligand 3,5-difluoro-4-hydroxybenzylidene-imidazolinone-2-oxime (DFHO) with nanomolar affinity. This binding event then activates fluorescence of DFHO, which when excited with light at a wavelength of 505 nm emits fluorescence at 545 nm. To enhance its stability to enable detection, the Corn aptamer is included within the middle of a tRNA scaffold sequence, which folds in such a way as to reduce RNA degradation [24]. Importantly for our purposes, the Corn aptamer system is both sufficiently photostable and transcribed at sufficient levels from the human U6 (hU6) Pol III promoter to enable the transcripts to be detected in cells using flow cytometry [24, 26, 27].

To employ these parts for the current study, we refined methods for quantifying RNA transcript levels in human cells [24]. We first sought to detect Pol III-driven transcription by expressing Corn aptamer-containing transcripts in the human embryonic kidney (HEK293FT) cell line (**Supplementary Fig. 1A**) [24]. Specifically, we transfected HEK293FT cells with the plasmid pAV-U6+27-tCORN, a plasmid construct containing in order, a human hU6 promoter, a 27 bp U6 leader sequence commonly included for optimal expression [27], the Corn aptamer fused to a tRNA scaffold, and a SV40 termination site [24]. In our experiments this initial design did not yield a signal that was significantly different than the background signal (**Supplementary Fig. 1B)**.

We hypothesized that this lack of signal may arise from inadequate transcript stability. Therefore, to boost the observable signal, we adapted the recently developed Tornado system which was designed to enhance the detectable signal from the Corn aptamer [26]. In the Tornado system, the Corn aptamer-tRNA scaffold is flanked by two twister ribozyme sequences (**Fig. 1A**). Following transcription, these self-cleaving ribozymes cleave the RNA in two locations, which allows the nuclear protein RtcB to ligate the free ends together, producing a circularized RNA containing the Corn aptamer. This circular RNA is protected from endogenous exonucleases, allowing Corn aptamer transcripts to accumulate to higher concentrations and thus conferring an enhancement in fluorescence (**Fig. 1A**) [26]. To utilize the Tornado system, we introduced a Reporter module into our constructs, consisting of an hU6 promoter, the same U6 leader sequence, a twister ribozyme, the Corn aptamer fused to a tRNA scaffold, a second twister ribozyme, and an SV40 termination site. When transfected into HEK293FT cells, this Tornado construct enabled robust detection of Pol III-driven transcription (**Fig. 1B**). We concluded that this Tornado-based system is well-suited for quantifying Pol III-driven expression and termination *in cellulo*.

### Poly-U Sequence Length Modulates Pol III Termination

We next sought to investigate how Pol III termination efficiency varies with the length of the poly-U sequence tract. To this end, we modified the Tornado reporter construct to include an additional Terminator module downstream of the U6 leader sequence and upstream of the Reporter module sequence elements (**Fig. 2A**). These Terminator modules incorporated a varying number of U nucleotides in the transcribed RNA. To study the effect of poly-U sequence length on termination independent of RNA structure, we included a computationally designed ‘linear’ sequence within the Terminator module, upstream of the poly-U sequence, which is predicted to lack any intramolecular RNA structures. Candidate linear sequences were designed using the Nucleic Acids PACKage (NUPACK) [28], and verified to be predicted to be singlestranded when included in a transcript alongside the entire Tornado Reporter module (**Fig. 2A, right; Methods; Supplementary Fig. 2**). Two different linear sequences were employed—Linear-1 and Linear-2—in order to evaluate whether any one specific choice of linear sequence contributes to termination.

**Figure 2:**
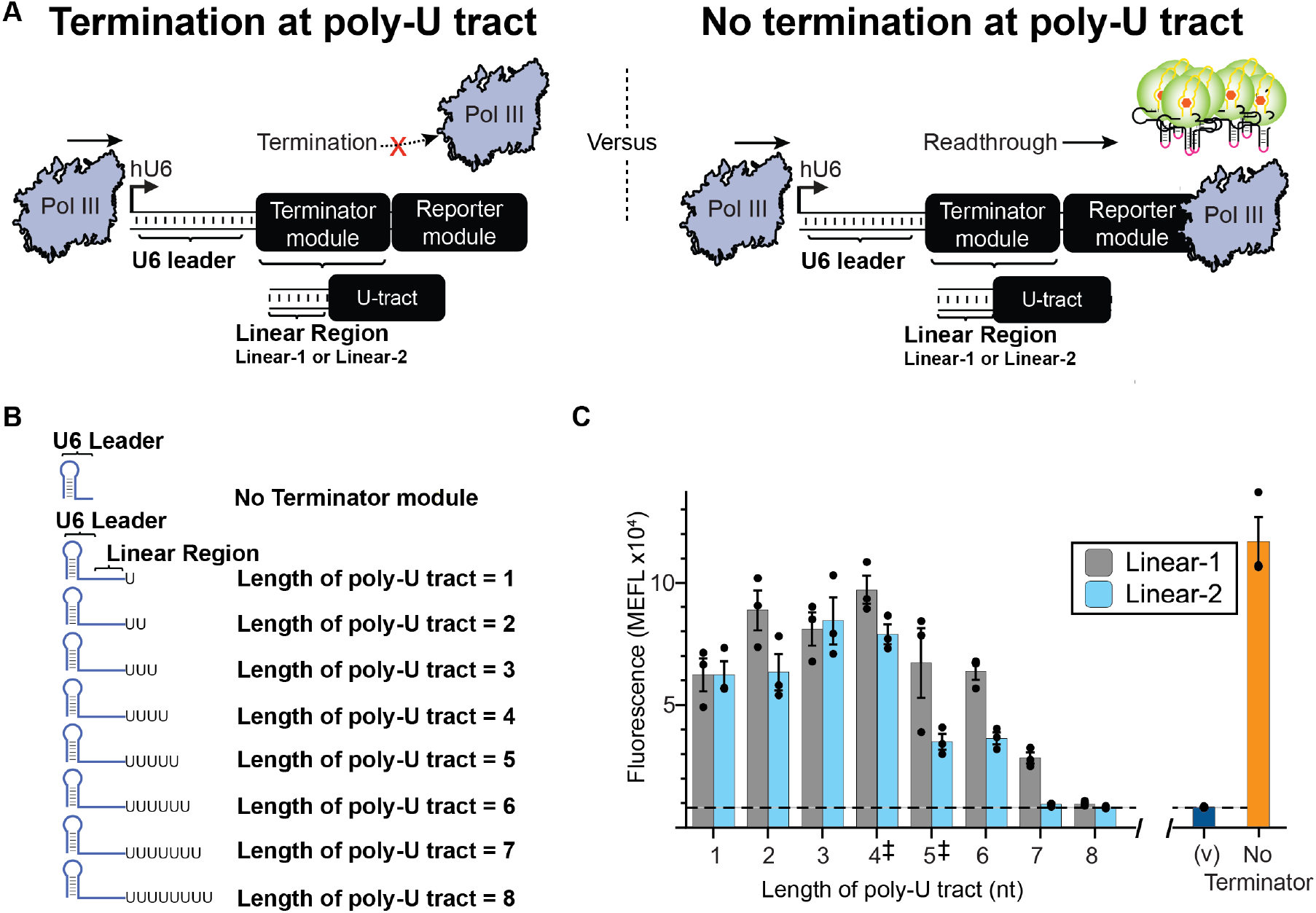
Varying poly-U tract length modulates termination of Pol III-driven transcription *in cellulo*. **A)** Assay design for quantifying relative termination efficiency as a function of poly-U tract sequence length. A ‘Terminator module’ was constructed by placing varying lengths of poly-U tract sequence downstream of RNA sequences designed to be completely linear—i.e., lacking secondary structure (Linear-1 or Linear-2). This combined Terminator module was placed immediately downstream of the U6 promoter and upstream of the Reporter module. In this system, the efficiency of the Terminator module is inversely related to the magnitude of the fluorescent output. **B)** Illustration of predicted RNA conformation which terminates at poly-U tracts of varying lengths. The U6 leader sequence is predicted to fold into a 5’ hairpin structure. **C)** Termination efficiency was experimentally quantified for constructs varying in poly-U length for two different linear sequences, with outputs compared to the fluorescence observed from cells transfected with a vector-only (no Reporter module) negative control (v), and a construct lacking a Terminator module. For both choices of linear sequence, poly-U tract lengths of 1-6 nt were found to be significantly different (p < 0.05) from the vector-only control using a one-tailed heteroscedastic Welch’s t-test followed by the Benjamini-Hochberg procedure with a false discovery rate cutoff of 0.05 (**Supplementary Table 2**). By this test, Linear-1 constructs with poly-U tracts of 7-8 nt also differed from the vector only control (p < 0.05), but Linear-2 constructs with tracts length of 7-8 nt did not. Colored bars represent the average of 3 biological replicates with individual points plotted as circles. Error bars represent the S.E.M. The dashed line represents the average of three replicates for the vector-only control (v) and the grey horizontal bar represents the S.E.M. of the vector-only control. ‡ denotes the two most commonly found poly-U tract lengths within the human genome.

Using this Terminator module approach, we evaluated the effect of poly-U tract lengths ranging from 1 to 8 nt on Pol III termination *in cellulo* (**Fig. 2B**). For both linear sequence contexts, we observed that past a certain length, increasing the poly-U tract length decreased reporter signal, indicating more efficient termination (**Fig. 2C**). Interestingly, within this assay, we actually observed a small increase in signal when comparing constructs containing poly-U tracts of length 4 nt to a length of 1 nt (**Supplementary Table 1**). This was surprising, as tract length of 4 uracils is the average length of all poly-U tracts in human Pol III-expressed genes [13]. We observed a trend of decreasing signal output only after poly-U tracts reached a size of 7 nt or greater. When comparing against the background signal from our vector-only control, only poly-U tracts of 7 or 8 uracils demonstrated no significant difference in observed signal (**Supplementary Table 2**). We speculated that if our model transcripts require longer poly-U tracts to achieve efficient termination than do endogenous Pol III-driven transcripts [13], perhaps other model transcript features could confer efficient termination with shorter poly-U tracts.

### RNA Structure Adjacent to the Poly-U Tract Enhances Pol III Termination

We next sought to investigate how upstream RNA structure might influence Pol III termination at poly-U tracts. We started by adapting our expression constructs to include a sequence that introduces a well-known secondary structural element by encoding a portion of the 5S ribosomal RNA (rRNA) hairpin. This 5S rRNA hairpin sequence was previously employed to investigate the impact of RNA secondary structure on Pol III termination using *in vitro* transcription assays [6]. In our investigation, we placed this sequence, predicted to fold into a 9 bp RNA hairpin structure with a 5 nt loop, immediately upstream of the poly-U tract (**Fig. 3A**). NUPACK analysis was then used to confirm that (i) the upstream linear region was still predicted to assume a single-stranded conformation, and (ii) no other competing RNA structures were predicted as a consequence of introducing the 5S rRNA hairpin (**Supplementary Fig. 2**). We first evaluated how adding this hairpin influences termination in the context of a 1 nt U tract. By itself, the single U did not cause significant termination (**Fig. 2B**), and no increase in termination (loss of reporter fluorescence) was observed due to the addition of the hairpin (**Fig. 3B**). Interestingly the addition of the hairpin with the 1nt U tract caused an increase in observed fluorescence in the Linear-2 sequence context. We next extended the poly-U tract to 4 nt, corresponding to the average poly-U length of Pol III transcripts in the human genome [13]. For these constructs, upon introduction of the hairpin, we observed significant decreases in fluorescence in comparison to all three of the previously tested conditions of (i) a poly-U tract of 1 uracil and no hairpin, (ii) a poly-U tract of 4 uracils and no hairpin, and (iii) a poly-U tract of 1 uracil and an adjacent hairpin. We observed this pattern for both choices of linear sequence. We conclude that within our setup, the presence of secondary structure in a nascent transcript enhances Pol III termination at a 4 nt poly-U tract.

### The Distance of the RNA Secondary Structure Impacts its Ability to Enhance Pol III Termination

We next investigated whether the position of the secondary structural element within the RNA transcript impacts its enhancement of termination efficiency. In the constructs analyzed in **Figure 3**, the hairpin was placed immediately adjacent to the poly-U track (i.e., a distance of 0 nt upstream). For comparison, we generated constructs in which the hairpin was instead placed on the other side of the linear sequence—at a distance of 10 nt upstream of the poly-U tract (**Supplementary Figure 3)**. We again employed NUPACK analysis to confirm that new transcripts were predicted to assume the expected conformations (**Supplementary Fig. 2**). For constructs with a 1 nt U tract, the addition of a hairpin at 10 nt upstream did not increase termination (**Supplementary Fig. 3B**). Rather, the addition of a hairpin in this position appeared to increase fluorescence **(Supplementary Table 1).** When the poly-U tract was extended to 4 nt for constructs with a hairpin 10 nt upstream of the poly-U tract, fluorescence decreased significantly in comparison to the other conditions tested (**Supplementary Fig. 3B**), indicating that even at this distance, secondary structure enhances Pol III termination. Similar trends were observed for both linear sequences. These data demonstrated that inclusion of a secondary structural element 10 nt upstream from the poly-U tract enhances Pol III termination in a similar fashion to that which occurs when the hairpin is directly adjacent to the poly-U tract.

**Figure 3:**
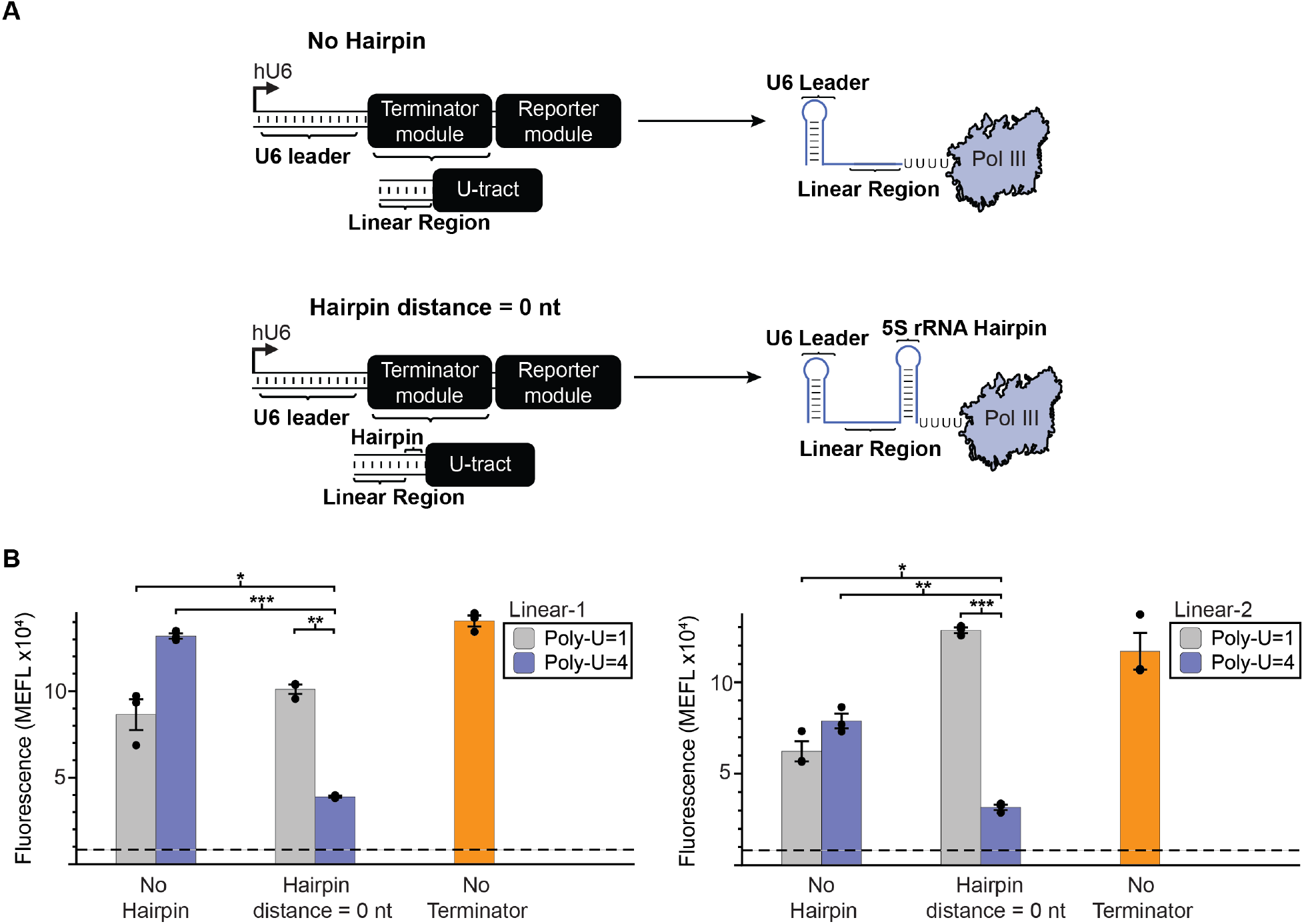
Upstream RNA secondary structure enhances termination as a function of poly-U tract length. **A)** A schematic depicting the positioning of an RNA secondary structure element immediately upstream of the poly-U tract within the Terminator module. The secondary structure utilized is a 23 nt portion of the 5S ribosomal RNA (rRNA) predicted to fold into a 9 bp hairpin following previous studies of Pol III termination *in vitro*. This structure was either omitted (No Hairpin) or positioned after the linear sequence and immediately upstream of the poly-U tract (Hairpin distance = 0 nt). **B)** Contribution of upstream RNA structure to termination was evaluated for both the Linear-1 (left) and Linear-2 (right) sequence contexts. Colored bars represent the average of 3 biological replicates with individual points plotted as circles. Error bars represent the S.E.M. The dashed line represents the average of three replicates for the vector-only control, and the grey horizontal bar represents the S.E.M. of the vector-only control., as reported in **Fig. 2C**. Statistical significance of the indicated comparisons (brackets) was measured using one-tailed heteroscedastic Welch’s t-tests followed by the Benjamini-Hochberg procedure with a false discovery rate cutoff of 0.05 (**Supplementary Table 2**). * = p < 0.05, ** = p< 0.01, *** = p < 0.001.

Interestingly, the above observation contrasts with the prokaryotic intrinsic termination mechanism, where the RNA hairpin must be placed immediately adjacent to the poly-U tract in order to confer effective termination [14]. We therefore sought to investigate how far upstream RNA structure can be placed and still enhance Pol III termination. To do so, we used NUPACK to design a new transcript with a longer linear sequence (Linear-3) that enables insertion of the RNA hairpin up to 20 nt upstream from the poly-U tract. First, we confirmed that Linear-3-based constructs exhibited the same patterns observed for Linear-1- and Linear-2-based constructs when the hairpin was placed 0 nt or 10 nt upstream from the poly-U tract (**Fig. 4B**). We then created a series of constructs by incrementally increasing the distance between the hairpin and the poly-U tract. Overall, we observed a general trend of increasing fluorescence (decreasing termination efficiency) with increasing distance between the hairpin and the poly-U tract; there was no observable impact on termination efficiency once this distance reached 20 nt (**Fig. 4C).**

**Figure 4:**
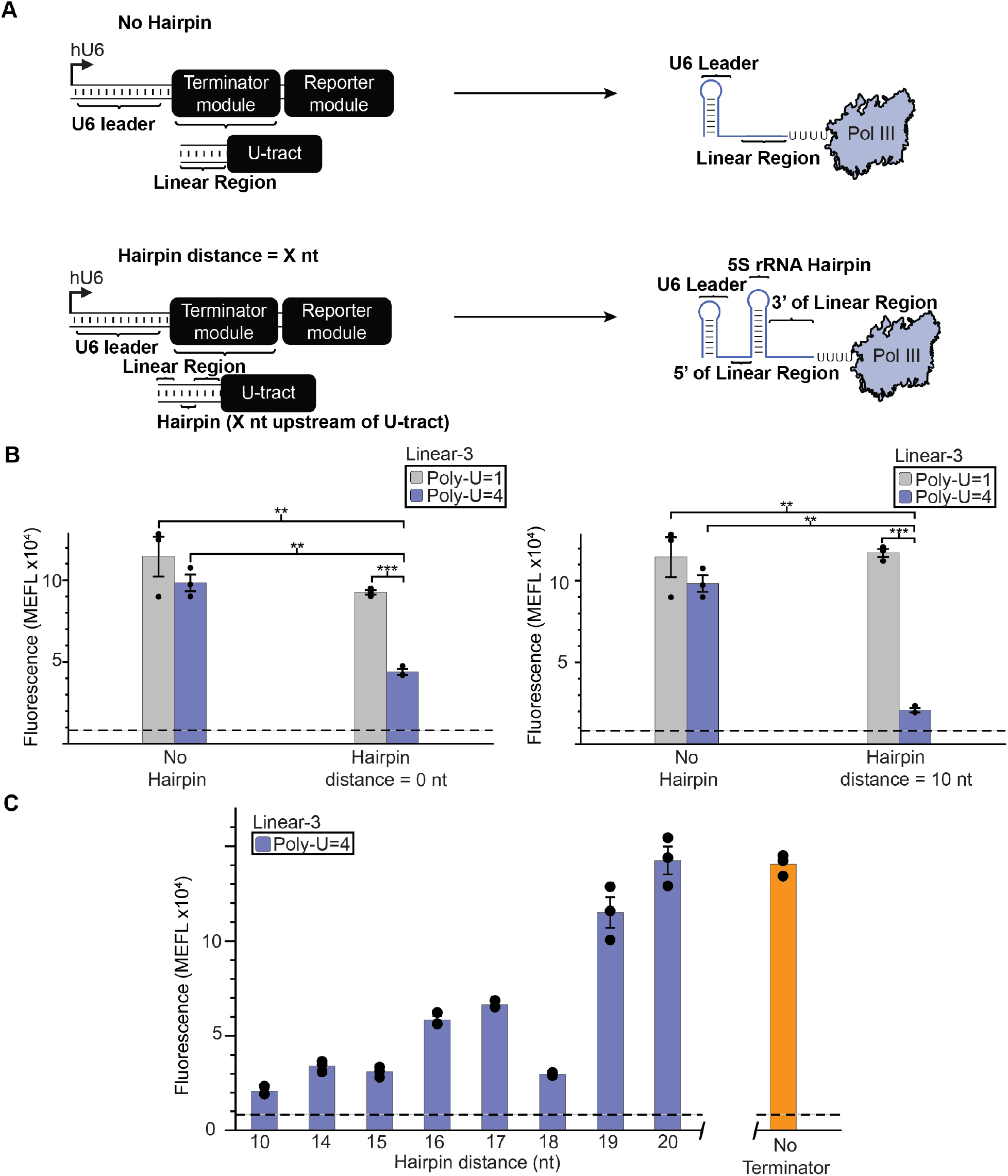
Distance of RNA structure from the poly-U tract impacts termination efficiency. **A)** A schematic depicting the positioning of secondary structure upstream of the poly-U tract. This structure was either omitted (No Hairpin) or positioned X nt upstream of the poly-U tract (where Hairpin distance = X nt). The secondary structure utilized is a 23 nt portion of the 5S ribosomal RNA (rRNA) predicted to fold into a 9 bp hairpin. In these constructs, the Linear-3 sequence was used to enable X to be up to 20 nt. **B)** Constructs based upon Linear-3 exhibit a pattern which is similar to those based upon Linear-1 and -2 for hairpin distances of 0 or 10 nt. Significance was measured using a one-tailed heteroscedastic Welch’s t-test followed by the Benjamini-Hochberg procedure with a false discovery rate cutoff of 0.05 (**Supplementary Table 2**). ** = p < 0.01, *** = p < 0.001. **C)** Moving the hairpin further upstream from the poly-U tract reduces termination efficiency. Observed fluorescence for hairpin distances 10, 14, 15, 16, 17, 18, 19 differed significantly from the no terminator control using the same statistical test (**Supplementary Table 2**), while observed fluorescence for a hairpin distance of 20 nt did not differ significantly from the No Terminator control. Colored bars represent the average of 3 biological replicates with individual points plotted as circles. Error bars represent the S.E.M. The dashed line represents the average of three replicates for the vector-only control (v) and the grey horizontal bar represents the S.E.M. of the vector-only control.

Overall, these data suggest that within the context of our assay, the location of RNA secondary structure and the length of the poly-U tract interact to modulate Pol III transcription termination efficiency in human cells.

## Discussion

In this study, we developed a means for quantitatively interrogating Pol III termination in human cells. We found that both poly-U sequence length, and the presence and position of an RNA hairpin structure can both influence Pol III termination. Specifically, we found that poly-U tracts alone can enhance transcription termination if they are at least 7 or 8 nts in length (**Fig. 2**). In addition, an RNA hairpin structure can enhance the termination when used in conjunction with shorter poly-U lengths (**Fig. 3**), although this effect is diminished the further away this hairpin structure is from the poly-U tract (**Fig. 4**). Notably, RNA structure by itself did not appear to cause termination (**Fig. 3**). This is an important advance in understanding, since previous studies evaluating the impact of multiple RNA sequence and structure elements on Pol III termination offered conflicting findings about the importance of these features [6, 7, 17, 18]. This study thus offers a potential resolution in supporting the interpretation that poly-U sequence and RNA structure are important for Pol III termination.

It is important to note the differences between this study and previous work. Notably, the previous studies utilized assays in yeast as well as *in vitro* transcription reactions with purified components. Therefore, the cellular context of these assays and our current work differ, and it is possible that our conclusions only specifically apply to human Pol III transcription termination.

Interestingly, the finding that both poly-U tract length and RNA secondary structure can enhance Pol III transcription is similar to the case of prokaryotic intrinsic termination [4]. Prokaryotic intrinsic termination occurs when the RNAP encounters a poly-U tract and changes from an elongation to paused state. During this pause, secondary structure encoded within the nascent RNA forms and acts to further destabilize the transcription complex resulting in transcription termination [16, 29]. In addition, the RNA secondary structure is often immediately adjacent to the poly-U tract in prokaryotes [14]. Notably, our findings for eukaryotic Pol III termination differ from the prokaryotic case in that the RNA structure still has an influence when not placed immediately adjacent to the poly-U tract. Potentially, this lack of spatial requirements may be due to the ability of Pol III to undergo extensive backtracking following interaction with the poly-U tract [6]. This backtracking of the polymerase may result in the repositioning of RNA secondary structure to be adjacent to the transcription complex, enabling termination. It is possible that the weaker hybridization forces between average length poly-U tracts and template result in a confirmational shift of the polymerase in a similar manner to what is seen during prokaryotic transcription termination [30, 31]. This shift may make the polymerase more sensitive to further destabilization forces, resulting in termination either from an upstream secondary structure or larger poly-U tracts. Further work will be needed to uncover the exact biomolecular interactions that are occurring during Pol III termination.

We also note that our synthetic biology approach for studying transcription termination using fluorescent RNA aptamers should be able to be used to study other features of Pol III termination. It would be of interest to assay the impacts of some notable factors such as the degree of upstream inter-nucleotide base stacking and the minimum free energy (MFE) of secondary structure [16]. Perhaps this work could also be utilized to perform functional enzymatic assays *in cellulo* to study how changes to the Pol III subunits, involved in termination, coordinate termination events with various nascent RNA sequence and structural elements [31]. We can envision testing an orthogonal Pol III mutant within our system following depletion of wild type Pol III that has been tagged with an inducible degradation systems [32]. As we currently do not expect any issues with adapting this assay for all culturable eukaryotic species, further testing may provide a complete model for Pol III termination across the eukaryotic domain.

This study adds to the growing body of knowledge of the RNA sequence and structural determinants of Pol III termination. Altogether, our system enables one to characterize termination within a context that may be most relevant for understanding the natural regulation of cellular processes which are known to impact both human development and disease [11, 33]. This could also be important from a biotechnology standpoint, as an increased understanding of Pol III termination may lead to forward-design of novel termination sequences that possess desired levels of termination in different genetic contexts, which could be useful for defining expression of genes useful in a range of biotechnology applications [15].

## Methods

### Design of RNA sequences

RNA sequences were designed, and structure prediction analysis performed using the Nucleic Acids PACKage [28]. RNA secondary structures were predicted from sequence utilizing the NUPACK online web portal in analysis mode. All folding queries were run under the RNA setting at 37°C using default parameters. Novel linear regions (i.e. RNA sequences predicted to not fold into secondary structures) were designed using the NUPACK web server in design mode by utilizing dot bracket notation of the desired length with the design feature. For example, to produce a hairpin with 5 base pairs and a loop of 4 nt, we input the notation (((((…))))), which NUPACK used to generate an RNA sequence predicted to fold into that structure. RNA sequence outputs were then inserted into the complete sequence construct to ascertain whether they were predicted to fold as designed in that context. Only sequences that were predicted to fold as designed were used in the study.

### Cloning

All cloning was done utilizing the Gibson assembly protocol or inverse PCR [34, 35]. All geneblocks and oligo primers were ordered from Integrated DNA Technologies. Assembled plasmids were transformed into and stored within NEB^®^ Turbo Competent Cells (**Supplementary Table 3**). All constructs were sequenced verified using Quintara Biosciences. All construct variants utilized in the main text were based off the construct pAV-U6+27-Tornado-Corn. The construct pAV-U6+27-tCORN, also from the Jaffrey lab, was obtained from Addgene (Addgene plasmid #106233) (**Supplementary Fig. 1**). A table of Addgene accession numbers for constructs utilized in this study, except for pAV-U6+27-Tornado-Corn, can be found in **Supplementary Table 4.** The construct pAV-U6+27-Tornado-Corn was received as a gift from Dr. Samie Jaffrey.

### Construct Preparation

Plasmids were transformed into NEB^®^ Turbo Competent Cells. Single colony forming units of each construct were then resuspended into 100 mL LB media with appropriate antibiotics and shook overnight at 37°C. Plasmids were harvested using Qiagen Midiprep Kits. Neutralized cell lysate was both spun down at 16xg for 30 min and the supernatant was then run through a Qiagen tip-100. Plasmids were resuspended in TE buffer at 200 ng/μL using a Thermo Scientific Nanodrop for quantification.

### Cell Culturing

HEK293FT cells (Life Technologies/Thermo) were maintained at 37°C and 5% CO2. Cells were cultured in DMEM (Gibco 31600-091) supplemented with 10% FBS, 6mM L-glutamine (2 mM from Gibco 31600-091 and 4 mM from additional Gibco 25030-081), penicillin (100 U/μL), and streptomycin (100 μg/mL) (Gibco 15140122).

### Transfection

HEK293FT cells were seeded at 1.5 x 10^5^ cells in 0.5 mL of supplemented DMEM media in 24 well plates. At 6-8 h post seeding, cells were transfected using calcium phosphate precipitation: DNA—200ng of construct (unless otherwise stated) (**Supplementary Fig. 4**) and 200ng of plasmid encoding blue fluorescent protein (BFP) as a transfection control—was mixed in H2O, and 2M CaCl2 was added to a final concentration of 0.3M CaCl2. The vector-only control was generated by transfecting with only 200ng of the plasmid encoding BFP. For all samples, transfected DNA mass was brought to a total of 800 ng/well though the addition of empty pcDNA™3.1(+) Mammalian Expression Vector (pcDNA) (ThermoFisher Scientific). This mixture was added dropwise to an equal-volume solution of 2x HEPES-Buffered Saline (280 mM NaCl, 0.5 M HEPES, 1.5 mM Na2HPO4) and gently pipetted up and down four times. After 2.5 min, the solution was mixed vigorously by pipetting ten times. 100 μL of this mixture was added dropwise to each well in a 24-well plate of cells. The next morning, the medium was aspirated and replaced with fresh medium.

### Sample Harvest

At 24-30 h post media change, cells were harvested for flow cytometry with 0.05% Trypsin EDTA (Thermo Fisher Scientific #25300120), incubating for 3 min at 37°C followed by quenching with phenol red-free DMEM. The resulting cell solution was added to 500 μL of flow buffer consisting of 1X phosphate-buffered saline (PBS), 4% FBS, 5 mM MgSO4 [24]. Cells were spun at 150xg for 5 minutes, supernatant was aspirated, and 200 μL of flow buffer supplemented to a final concentration of 5 μM DFHO dye (TOCRIS Bioscience) was added prior to analysis by flow cytometry.

### Analytical Flow Cytometry

Flow cytometry was performed using on a BD LSR Fortessa Special Order Research Product. Approximately 3,000-6,000 single transfected cells were analyzed per sample, using BFP as the transfection control. BD LSR Fortessa settings used were as follows: BFP was collected in the Pacific Blue channel (405 nm excitation, 450/50 nm filter) and DFHO dye signal was collected in the FITC channel (488 nm excitation, 505 LP and 530/30 nm filter). Samples were analyzed using FlowJo v10 software (FlowJo, LLC). Fluorescence data were compensated for spectral bleed-through, the HEK293FT cell population was identified by SSC-A vs. FSC-A gating, and single cells were identified by FSC-A vs. FSC-H gating (**Supplementary Fig. 5**). To distinguish between transfected and non-transfected cells, a control sample of cells was transfected with empty vector only (pcDNA). This empty vector control was used to identify cells that were positive for the constitutive fluorescent protein (BFP) used as a transfection control in all other samples. The gate was drawn to include no more than 1% of cells in the empty vector control. Constructs with reporters for the respective FITC and BFP fluorescence channels were analyzed and compensation was applied to account for spectral overlap (**Supplementary Fig. 5**)

After gating for transfection, the intensity of the DFHO dye signal was quantified as Mean Fluorescence Intensity (MFI) by taking the geometric mean of fluorescence intensity in the FITC channel within each transfected cell population. MFI was then converted to Mean Equivalent of Fluorescien (MEFL) using UltraRainbow Calibration Particles (Spherotech URCP-100-2H), which were incorporated as part of each individual experiment (**Supplementary Fig. 6**). The bead population was identified by FSC-A vs. SSC-A gating, and 9 bead subpopulations were identified through two fluorescent channels. MEFL values corresponding to each subpopulation were supplied by the manufacturer and a calibration curve was generated for the experimentally determined MFI vs. the manufacturer specified MEFLs. A linear regression was performed with the constraint that 0 MFI equals 0 MEFL, and the slope from the regression was used to convert MFI to MEFL for each cellular population.

Finally, we confirmed that reporter output did not vary significantly across the course of a representative 1.5 h flow cytometry data collection experiment (**Supplementary Fig. 7**), ruling out potential artifacts due to the order in which samples were analyzed.

### Data Analysis

The mean fluorescence intensity (MFI) described above of the singlecell, transfected population was calculated and exported for further analysis. To calculate signal, the FITC channel MFI was averaged across three biological replicates. A vector-only control sample transfected with the BFP transfection control and empty vector (pcDNA) was treated with 5 μM DFHO dye, averaged across three biological replicates, and used to measure background signal. Statistical significance of measured fluorescence differences between specified cell populations was measured by utilizing a one-tailed heteroscedastic Welch’s t-test either alone, or followed by the Benjamini-Hochberg procedure [36]. Data and statistical analysis were performed using Excel (Microsoft). (**Supplementary Table 1 & Supplementary Table 2**).

## Data Availability

All processed analytical flow cytometry data, as well as calculations of averages, S.E.M. and statistical comparisons, was deposited within Northwestern’s open access arch database (https://arch.library.northwestern.edu/). The data can be accessed at https://doi.org/10.21985/n2-9je1-6c04

## Author Contributions

M.S.V. conceived the project. M.S.V., W.K.C., T.B.D., J.N.L. & J.B.L curated the data. M.S.V., W.K.C., & T.B.D carried out the formal analysis. J.N.L & J.B.L. acquired the funds. M.S.V., W.K.C., & T.B.D carried out the investigations. M.S.V., W.K.C., & T.B.D devised the methodology. M.S.V., & J.N.L., and J.B.L. administered the project. M.S.V., W.K.C., & T.B.D acquired the resources. M.S.V., J.N.L, and J.B.L. supervised the project. M.S.V., W.K.C., T.B.D., J.N.L. & J.B.L validated the project. M.S.V. visualized the project. M.S.V. & T.B.D. wrote the original draft. M.S.V., W.K.C., T.B.D., J.N.L. & J.B.L reviewed and edited the manuscript.

## Acknowledgment

We wish to thank Dr. Samie Jaffrey for sharing a stock of the original Tornado construct bearing the Corn aptamer. We wish to acknowledge Michael Sumner for initial conversations around the ideas that led to this work. We would also like to acknowledge Adam Silverman for critical reading of this manuscript. M.S.V and W.K.C. were supported in part by Northwestern University’s Graduate School Cluster in Biotechnology, System, and Synthetic Biology, which is affiliated with the Biotechnology Training Program. W.K.C. was supported in part by the National Institutes of Health Training Grant (T32GM008449) through Northwestern University’s Biotechnology Training Program. This work was also supported by the Northwestern University – Flow Cytometry Core Facility supported by Cancer Center Support Grant (NCI CA060553). This work was supported in part by the National Institute of Biomedical Imaging and Bioengineering (grant no. 1R01EB026510 to J.N.L.), an NSF CAREER (grant no. 1452441 to J.B.L.), and through the National Institute for General Medicine Sciences (grant no. R01GM130901 to J.B.L).

