## Supplementary Information for "RNA sequence and structure determinants of Pol III transcriptional termination in human cells"

**Table of Contents**


- Supplementary Figure 1:Evaluating Corn aptamer detection in the context of linear RNA transcripts
- Supplementary Figure 2: Predicted secondary structure of representative transcripts.
- Supplementary Figure 3: RNA secondary structure 10 nt upstream enhance termination with some poly-U tracts.
- Supplementary Figure 4: Influence of Tornado construct DNA dose on output magnitude.
- Supplementary Figure 5: Flow cytometry gating workflow.
- Supplementary Figure 6: Flow cytometry fluorescence intensity calibration to absolute units with UltraRainbow beads.
- Supplementary Figure 7: Evaluation of signal variation of the positive control Tornado construct within an experiment.

- Supplementary Table 1: All statistical comparisons not utilizing multiple hypothesis correction procedures.
- Supplementary Table 2: Statistical comparisons using the Benjamini-Hochberg procedure.
- Supplementary Table 3: Sequences of DNA parts and constructs.
- Supplementary Table 4: Addgene ID # for plasmids generated for this study.

- References cited in Supplementary Information.

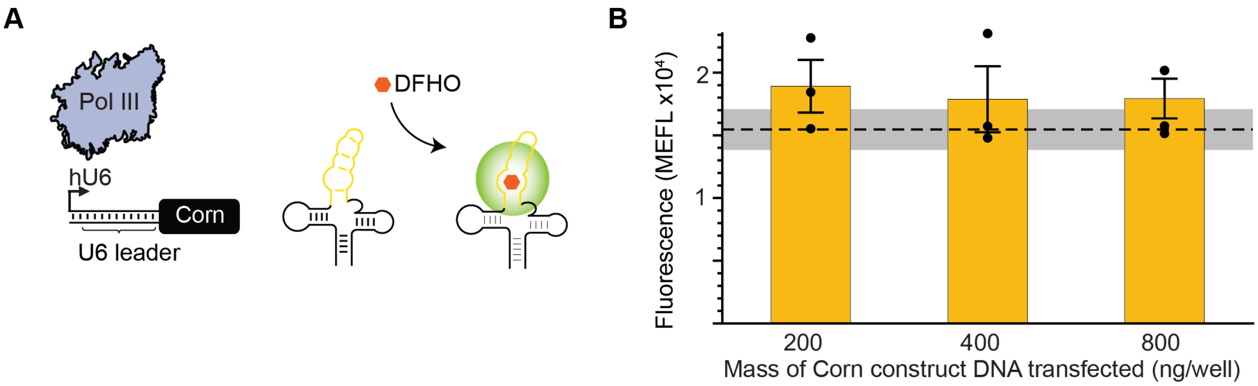


**Supplementary Figure 1: Evaluating Corn aptamer detection in the context of linear RNA transcripts. A)** A schematic overview of an expression cassette for evaluating observed signal from the fluorescent aptamer Corn expressed as a Pol III-driven transcript. The system comprises a human U6 promoter driving transcription of a Corn aptamer (yellow line) placed within a tRNA scaffold. **B)** Various doses of a construct bearing this expression cassette were transfected into HEK293FT cells, holding total DNA dose constant using empty vector DNA. Then, 48 h later, DFHO was added to the culture medium, and cells were harvested for analysis by flow cytometry. The resulting signal was compared to that observed for a vector-only control. None of these cases differed substantially from the vector-only control as measured by a 1-sided heteroscedastic Welch's t-test followed by the Benjamini-Hochberg procedure with a false discovery rate cutoff of 0.05 (**Supplementary Table 2**). Colored bars represent the average of 3 biological replicates with individual points plotted as circles. Error bars represent the S.E.M. The dashed line represents the average of three replicates for the vector-only control (v) and the grey horizontal bar represents the S.E.M. of the vector-only control.

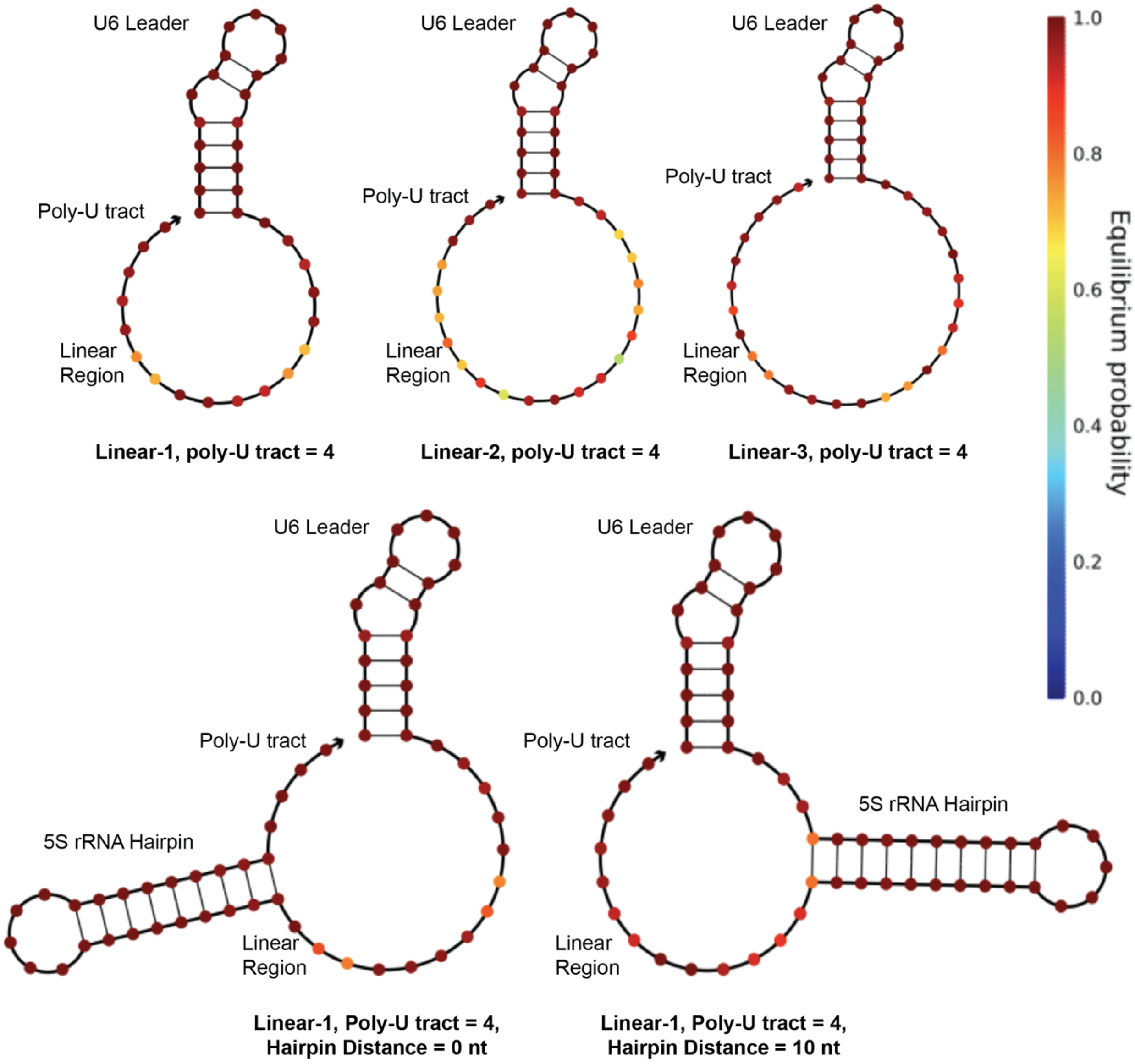


**Supplementary Figure 2: Predicted secondary structure of representative transcripts.** NUPACK [1] secondary structure analysis was used to predict the structural conformations of RNA sequences included in this study. Equilibrium folding analysis at 37 ºC using default NUPACK parameters used sequences beginning at the transcription start site (beginning of the U6 leader sequence) through the end of the terminator module. Shown here are transcripts possessing a 4 nt poly-U tract. Colored positions (colorbar) indicate the probability that nucleotides are in their indicated structural state (paired or unpaired) over the entire ensemble of possible structures. These examples illustrate sequences predicted to have a high probability of exhibiting the desired structure (e.g., single-stranded for linear regions, correctly base-paired within the U6 leader and 5S ribosomal RNA hairpin).

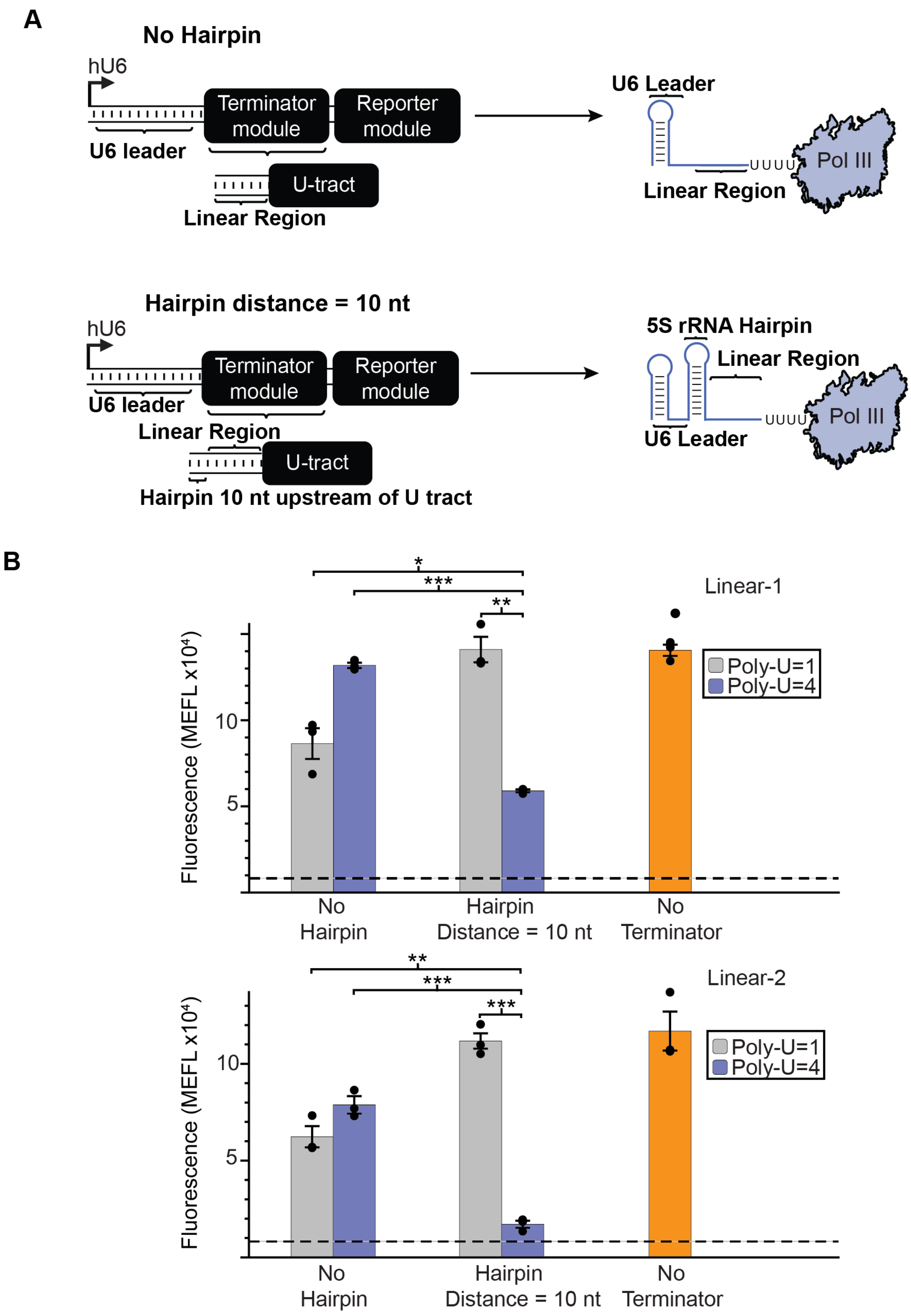

A
No Hairpin
hU6
Terminator module
Reporter module
U6 leader
U-tract
Linear Region
U6 Leader
UUUU
Pol III
Linear Region
Hairpin distance = 10 nt
hU6
Terminator module
Reporter module
U6 leader
Linear Region
U-tract
Hairpin 10 nt upstream of U tract
5S rRNA Hairpin
Linear Region
UUUU
Pol III
U6 Leader
B
Linear-1
*
***
**
Fluorescence (MEFL x10⁴)
10
5
Poly-U=1
Poly-U=4
No Hairpin
Hairpin Distance = 10 nt
No Terminator
Linear-2
**
***
***
Fluorescence (MEFL x10⁴)
10
5
Poly-U=1
Poly-U=4
No Hairpin
Hairpin Distance = 10 nt
No Terminator

**Supplementary Figure 3: RNA secondary structure 10 nt upstream of some poly-U tracts enhances termination. A)** A schematic depicting the positioning of secondary structure upstream of the poly-U tract. This structure was either omitted (No Hairpin) or placed 10 nt upstream of the poly-U tract (Hairpin distance = 10 nt). The secondary structure utilized is a 23 nt portion of the 5S ribosomal RNA (rRNA) predicted to fold into a 9 bp hairpin following previous studies of Pol III termination *in vitro* [2]. **B)** The two different configurations were included upstream of poly-U tracts of length 1 and 4 nt for both the Linear-1 and Linear-2 sequence contexts. Colored bars represent the average of 3 biological replicates with individual points plotted as circles. Error bars represent the S.E.M. The dashed line represents the average of three replicates for the vector-only control (v) and the grey horizontal bar represents the S.E.M. of the vector-only control. Statistical significance of the indicated comparisons (brackets) was measured using a one-tailed heteroscedastic Welch's t-test followed by the Benjamini-Hochberg procedure with a false discovery rate cutoff of 0.05 (**Supplementary Table 2**). * = $p < 0.05$, ** = $p < 0.01$, *** = $p < 0.001$.

**A** **B**

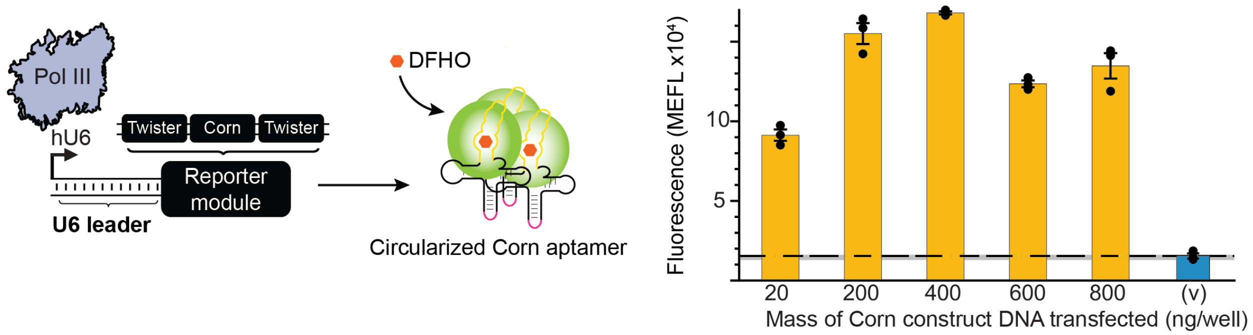


**Supplementary Figure 4: Influence of Tornado construct DNA dose on output magnitude. A)** Schematic of the tornado reporter module system from Fig. 1A. **B)** This construct was transfected into HEK293FT cells at various concentrations. For all samples, total DNA dose was held constant at 800 ng/well by adding in the required amount of empty vector DNA. The resulting signal was compared against the background fluorescence of the system measured by assaying cells transfected only with empty vector DNA (v). Colored bars represent the average of 3 biological replicates with individual points plotted as circles. Error bars represent the S.E.M. The dashed line represents the average of three replicates for the vector-only control (v), and the grey horizontal bar represents the S.E.M. of the vector-only control.

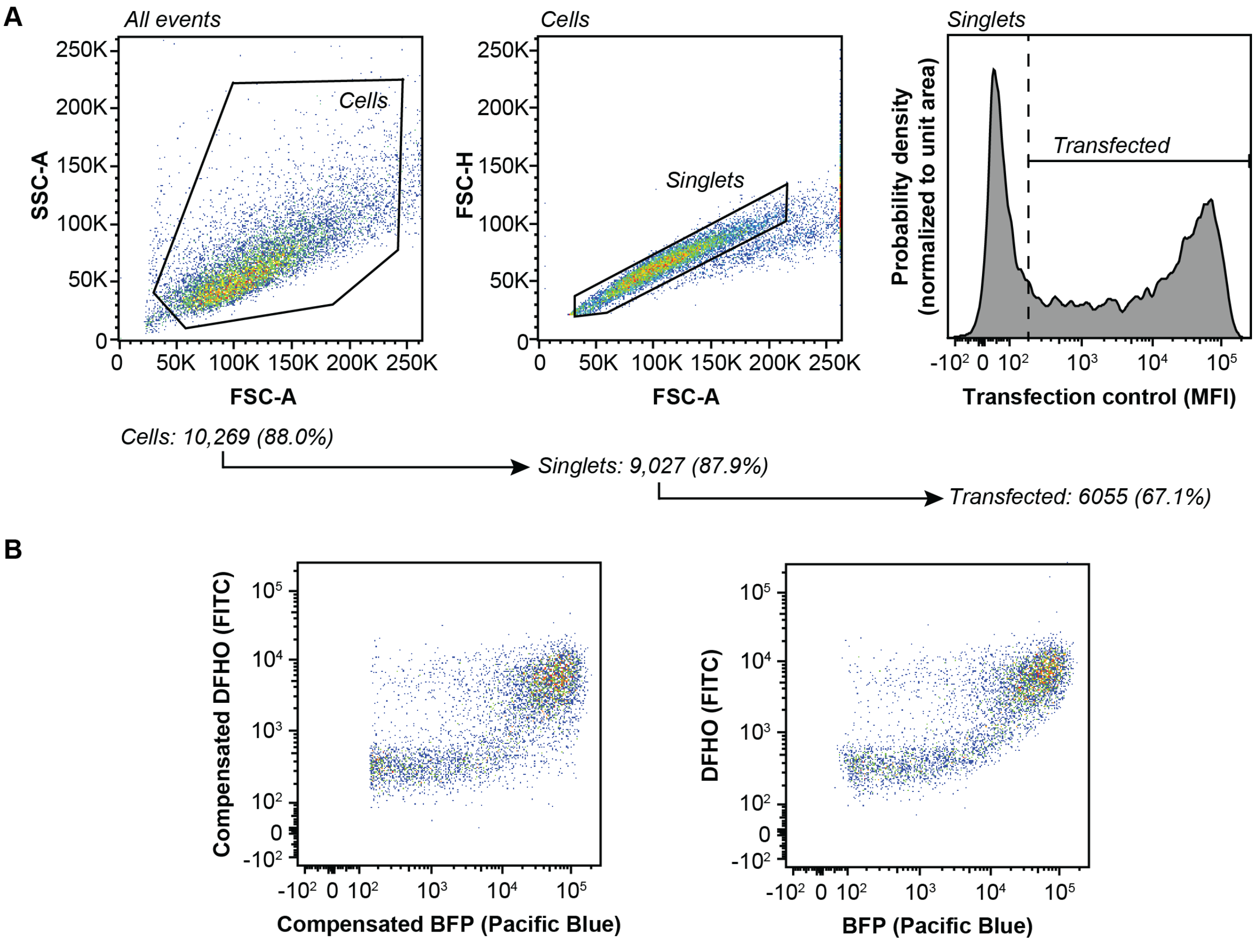


**Supplementary Figure 5: Flow cytometry gating workflow. A)** The plot shows cells transfected with empty vector only (pcDNA) that did not contain TORNADO expression cassettes and were treated with DFHO dye to illustrate the flow cytometry gating strategy used to identify single transfected cells. HEK293FT cells were identified by the FSC-A vs SSC-A profile (left). From this population, single cells were identified by the FSC-A vs. FSC-H profile (middle). Mean fluorescence intensity (MFI) of each cell was then collected and plotted as a distribution (right). The transfected population was defined as all single cells with a transfection control signal greater than the sample of single cells transfected with empty vector only, encompassing no more than 1% of the empty vector only transfected population. B) This figure illustrates the application of compensation to minimize spectral overlap between FITC and BFP signals. In this sample (only BFP+ cells are shown), all cells were transfected with constructs transcribing the Corn aptamer without an upstream Terminator module, and cells were treated with DFHO; cells receiving small amounts of these transfected plasmids (low BFP transfection control signal) are also low in FITC signal, as expected. Application of compensation (left) enables one to isolate the FITC and BFP signals, which in this experiment causes better separation of the various subpopulations compared to the uncompensated (right) version of these data.

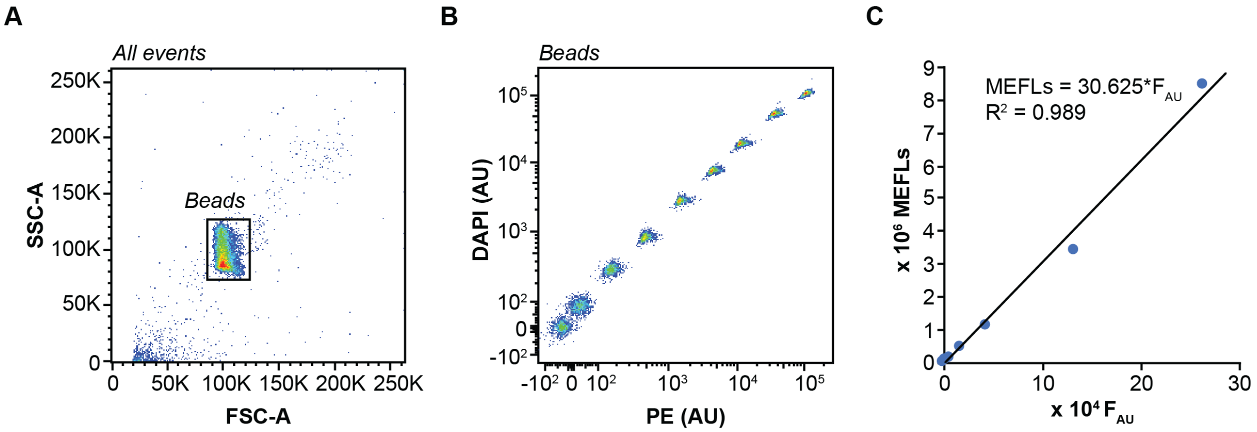


**Supplementary Figure 6: Flow cytometry fluorescence intensity calibration to absolute units with UltraRainbow beads. A)** The bead population was identified based on the FSC-A vs. SSC-A profile. **B)** Two fluorescent channels, other than the channels of interest for the experiment, were used to identify the 9 beads population. **C)** The mean fluorescence intensity (MFI) of each population in the FITC channel (used for DFHO dye signal) in arbitrary unites ($F_{AU}$) was recorded and plotted against manufacturer provided values for molecules of equivalent fluorescein (MEFL) per bead peak for each population. A linear regression with y-intercept set to zero was used to create a calibration curve. This curve was used to convert exported MFI values to absolute units using the multiplier obtained from the regression for each characterized cell population that was gated for transfection as in **Supplementary Figure 5**.

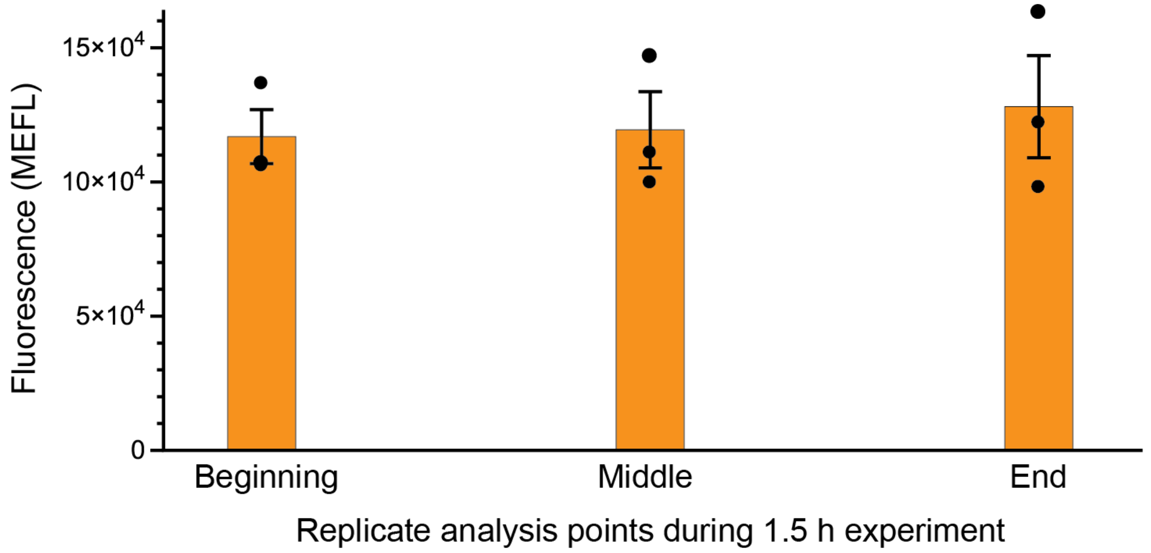


**Supplementary Figure 7: Evaluation of signal variation of the positive control Tornado construct within an experiment.** To identify potential sources of error and avoid misinterpretation, we evaluated whether reporter signal varies across the course of a single analytical flow cytometry data collection experiment. Within a 1.5 h long experiment, three samples (three biological replicate each) of cells transfected to express the Tornado construct (**Fig. 1A**) were assayed, at the beginning, middle, and end of the experiment, respectively. No statistical difference was observed between these points using a 2-tailed heteroscedastic Welch's t-test with no Benjamini-Hochberg correction (**Supplementary Table 1**). Colored bars represent the average of 3 biological replicates with individual points plotted as circles. Error bars represent the S.E.M.

**Supplementary Table 1: All statistical comparisons not utilizing multiple hypothesis correction procedures.** Each figure utilizing a Welch's t-test without a Benjamini-Hochberg is tabulated to demonstrate the statistical significance between tested points.

| Figure | Comparison | # sided t-test | p-value |
|---|---|---|---|
| Fig. 1B | Tornado System vs Vector-only control | 1 | 0.00066645 |
| Fig. 2C | Linear-1, PolyU=1 vs Linear-1, PolyU=4 | 1 | 0.00907346 |
| Fig. 2C | Linear-2, PolyU=1 vs Linear-2, PolyU=4 | 1 | 0.03881797 |
| Supp. Fig. 3B | Linear-1, PolyU=1 vs Linear-1, PolyU=1, Hairpin Distance =10nt | 1 | 0.00502829 |
| Supp. Fig. 3B | Linear-2, PolyU=1 vs Linear-2, PolyU=1, Hairpin Distance =10nt | 1 | 0.00127521 |
| Supp. Fig 7 | Beginning vs (Middle & End) | 2 | 0.65795305 |
| Supp. Fig 7 | Middle vs (Beginning & End) | 2 | 0.73657755 |
| Supp. Fig 7 | End vs (Beginning & End) | 2 | 0.66684677 |

**Supplementary Table 2: Statistical comparisons using the Benjamini-Hochberg procedure.** Multiple statistical comparisons utilizing a 1-sided heteroscedastic Welch's t-test were corrected following the Benjamini-Hochberg procedure in order to control the false discover rate (FDR) [3]. For each figure where this is used, rank order t-tests are given along with the comparisons where the null hypothesis (that the comparisons are equal) is rejected (TRUE) or not rejected (FALSE) for three different FDR cutoffs.

| Figure | Ranked Comparison | Ranked t-test | Null hypothesis rejected | | |
|---|---|---|---|---|---|
| Fig. 2C | vs Vector-only Control (v) | | FDR=0.05 | FDR=0.01 | FDR=0.001 |
| | Linear-2, PolyU(4) | 1.53E-03 | TRUE | FALSE | FALSE |
| | Linear-1, PolyU(6) | 1.94E-03 | TRUE | FALSE | FALSE |
| | Linear-1, PolyU(4) | 2.14E-03 | TRUE | FALSE | FALSE |
| | Linear-2, PolyU(6) | 3.45E-03 | TRUE | FALSE | FALSE |
| | Linear-1, PolyU(3) | 4.25E-03 | TRUE | FALSE | FALSE |
| | Linear-2, PolyU(1) | 5.02E-03 | TRUE | FALSE | FALSE |
| | Linear-1, PolyU(2) | 5.14E-03 | TRUE | FALSE | FALSE |
| | Linear-1, PolyU(7) | 5.58E-03 | TRUE | FALSE | FALSE |
| | Linear-2, PolyU(5) | 6.74E-03 | TRUE | FALSE | FALSE |
| | Linear-1, PolyU(1) | 7.51E-03 | TRUE | FALSE | FALSE |
| | Linear-2, PolyU(3) | 7.90E-03 | TRUE | FALSE | FALSE |
| | Linear-2, PolyU(2) | 8.81E-03 | TRUE | FALSE | FALSE |
| | Linear-1, PolyU(5) | 2.68E-02 | TRUE | FALSE | FALSE |
| | Linear-1, PolyU(8) | 3.67E-02 | TRUE | FALSE | FALSE |
| | Linear-2, PolyU(7) | 6.10E-02 | FALSE | FALSE | FALSE |
| | Linear-2, PolyU(8) | 4.08E-01 | FALSE | FALSE | FALSE |
| Fig. 3B | vs Linear-1, PolyU(4), Hairpin Distance = 0nt | | | | |
| | Linear-1, PolyU(4), No Hairpin | 3.98E-05 | TRUE | TRUE | TRUE |
| | Linear-1, PolyU(1), Hairpin Distance 0nt | 7.44E-04 | TRUE | TRUE | FALSE |
| | Linear-1, PolyU(1), No Hairpin | 1.67E-02 | TRUE | FALSE | FALSE |
| Fig. 3B | vs Linear-2, PolyU(4), Hairpin Distance = 0nt | | | | |
| | Linear-2, PolyU(1), Hairpin Distance 0nt | 7.01E-07 | TRUE | TRUE | TRUE |
| | Linear-2, PolyU(4), No Hairpin | 1.58E-03 | TRUE | TRUE | FALSE |

| | | | | | |
|---|---|---|---|---|---|
| | Linear-2, PolyU(1), No Hairpin | 1.21E-02 | TRUE | FALSE | FALSE |
| Fig. 4B | vs Linear-3, PolyU(4), Hairpin Distance 0nt | | | | |
| | Linear-3, PolyU(1), Hairpin Distance 0nt | 2.45E-05 | TRUE | TRUE | TRUE |
| | Linear-3, PolyU(4), No Hairpin | 2.19E-03 | TRUE | TRUE | FALSE |
| | Linear-3, PolyU(1), No Hairpin | 1.36E-02 | TRUE | TRUE | FALSE |
| Fig. 4B | vs Linear-3, PolyU(4), Hairpin Distance 10nt | | | | |
| | Linear-3, PolyU(1), Hairpin Distance 10nt | 3.06E-05 | TRUE | TRUE | TRUE |
| | Linear-3, PolyU(4), No Hairpin | 1.29E-03 | TRUE | TRUE | FALSE |
| | Linear-3, PolyU(1), No Hairpin | 7.99E-03 | TRUE | TRUE | FALSE |
| Fig. 4C | vs No Terminator | | | | |
| | Linear-3, PolyU(4), Hairpin Distance 15nt | 4.53E-05 | TRUE | TRUE | TRUE |
| | Linear-3, PolyU(4), Hairpin Distance 16nt | 5.18E-05 | TRUE | TRUE | TRUE |
| | Linear-3, PolyU(4), Hairpin Distance 10nt | 6.26E-05 | TRUE | TRUE | TRUE |
| | Linear-3, PolyU(4), Hairpin Distance 14nt | 6.26E-05 | TRUE | TRUE | TRUE |
| | Linear-3, PolyU(4), Hairpin Distance 18nt | 3.27E-04 | TRUE | TRUE | TRUE |
| | Linear-3, PolyU(4), Hairpin Distance 17nt | 3.36E-04 | TRUE | TRUE | TRUE |
| | Linear-3, PolyU(4), Hairpin Distance 19nt | 3.62E-02 | TRUE | FALSE | FALSE |
| | Linear-3, PolyU(4), Hairpin Distance 20nt | 4.14E-01 | FALSE | FALSE | FALSE |
| Fig. S1 | vs Vector-only control (v) | | | | |
| | Corn [200ng] | 1.34E-01 | FALSE | FALSE | FALSE |
| | Corn [400ng] | 3.72E-03 | FALSE | FALSE | FALSE |
| | Corn [800ng] | 4.53E-02 | FALSE | FALSE | FALSE |
| Fig. S4 | Linear-1, PolyU(4) , Hairpin Distance 10nt | | | | |
| | Linear-1, PolyU(4), No Hairpin | 1.34E-01 | TRUE | TRUE | TRUE |
| | Linear-1, PolyU(1), Hairpin Distance 10nt | 2.43E-01 | TRUE | TRUE | FALSE |
| | Linear-1, PolyU(1), No Hairpin | 2.64E-01 | TRUE | FALSE | FALSE |
| Fig. S4 | Linear-2, PolyU(4), Hairpin Distance 10nt | | | | |
| | Linear-2, PolyU(1), Hairpin Distance 10nt | 3.12E-04 | TRUE | TRUE | TRUE |
| | Linear-2, PolyU(4), No Hairpin | 5.25E-04 | TRUE | TRUE | TRUE |
| | Linear-2, PolyU(1), No Hairpin | 4.36E-03 | TRUE | TRUE | FALSE |

**Supplementary Table 3: Sequences of DNA parts and constructs.**

| Part/Construct | Sequence |
|---|---|
| U6 Promoter | gagggcctatttcccatgattccttcatatttgcatatacgatacaaggctgttagagagataatt agaattaatttgactgtaaacacaaagatattagtacaaaatacgtgacgtagaaagtaata atttcttgggtagtttgcagttttaaaattatgttttaaaatggactatcatatgcttaccgtaacttga aagtatttcgatttcttggctttatatatcttgtggaaaggac |
| 27 bp Leader | gtgctcgcttcggcagcacatatactag |
| Linear 1: | ttggc |
| Linear 2: | tgattgatg |
| Linear 3: | cagccaactccaa |
| 5S rRNA Hairpin | ggttgcggctttctgctgcaatc |
| Corn/tRNA scaffold | caggcccggatagctcagtcggtagagcagcggccgcgcgattcgaggaaggaggtctg aggaggtcactgaatcgcgcggccgcgggtccagggttcaagtccctgttcgggcctgccat cagtc |
| 5' Twister ribozyme | ggccatcagtcgccggtcccaagcccggataaaatgggagggggcgggaaaccgcctaa ccatgccgactgatgg |
| 3' Twister ribozyme | ggcgtggactgtagaacactgccaatgccggtcccaagcccggataaaagtggagggtac agtccacgc |
| SV40 Terminator | aacttgtttattgcagcttataatggttacaaataaagcaatagcatcacaaatttcacaaataa agcatttttttcactgcattctagttgtggtttgtccaaactcatcaatgtatctta |
| Construct: Linear:1 with a poly-U tract of 4 and secondary structure immediately adjacent (Fig. 2B). | gagggcctatttcccatgattccttcatatttgcatatacgatacaaggctgttagagagataatt agaattaatttgactgtaaacacaaagatattagtacaaaatacgtgacgtagaaagtaata atttcttgggtagtttgcagttttaaaattatgttttaaaatggactatcatatgcttaccgtaacttga aagtatttcgatttcttggctttatatatcttgtggaaaggacgaaacaccgtgctcgcttcggca gcacatatactagttggcggttgcggctttctgctgcaatcttttcgacggccatcagtcgccggt cccaagcccggataaaatgggagggggcgggaaaccgcctaaccatgccgactgatggc aggcccggatagctcagtcggtagagcagcggccgcgcgattcgaggaaggaggtctga ggaggtcactgaatcgcgcggccgcgggtccagggttcaagtccctgttcgggcctgccatc agtcggcgtggactgtagaacactgccaatgccggtcccaagcccggataaaagtggagg gtacagtccacgctctagagcggacttcggtccgctttttactaggacctgcaggcatgcaag cttgacgtcggttaccgatatccatatggcggccgcatcgatctcgagccgcggactagtaac ttgtttattgcagcttataatggttacaaataaagcaatagcatcacaaatttcacaaataaagc atttttttcactgcattctagttgtggtttgtccaaactcatcaatgtatctta |

**Supplementary Table 4: Addgene ID # for plasmids generated for this study.** The following constructs are or will be made available on Addgene.org.

| Addgene ID # | Construct Name |
|---|---|
| 106233 | pAV-U6+27-tCORN |
| 159479 | pAV-U6+27-Linear-1_PolyU(1)_ |
| 159480 | pAV-U6+27-Linear-1_PolyU(2)_ |
| 159481 | pAV-U6+27-Linear-1_PolyU(3)_ |
| 159482 | pAV-U6+27-Linear-1_PolyU(4)_Tornado-Corn_ |
| 159483 | pAV-U6+27-Linear-1_PolyU(5)_Tornado-Corn_ |
| 159484 | pAV-U6+27-Linear-1_PolyU(6)_Tornado-Corn_ |
| 159485 | pAV-U6+27-Linear-1_PolyU(7)_Tornado-Corn_ |
| 159486 | pAV-U6+27-Linear-1_PolyU(8)_Tornado-Corn_ |
| 159487 | pAV-U6+27-Linear-2_PolyU(1)_Tornado-Corn_ |
| 159488 | pAV-U6+27-Linear-2_PolyU(2)_Tornado-Corn_ |
| 159489 | pAV-U6+27-Linear-2_PolyU(3)_Tornado-Corn_ |
| 159490 | pAV-U6+27-Linear-2_PolyU(4)_Tornado-Corn_ |
| 159491 | pAV-U6+27-Linear-2_PolyU(5)_Tornado-Corn_ |
| 159492 | pAV-U6+27-Linear-2_PolyU(6)_Tornado-Corn_ |
| 159493 | pAV-U6+27-Linear-2_PolyU(7)_Tornado-Corn_ |
| 159494 | pAV-U6+27-Linear-2_PolyU(8)_Tornado-Corn_ |
| 159495 | pAV-U6+27-Linear-1_PolyU(1)_HairpinDistance(0)nts_Tornado-Corn_ |
| 159496 | pAV-U6+27-Linear-1_PolyU(4)_HairpinDistance(0)_Tornado-Corn_ |
| 159497 | pAV-U6+27-Linear-1_PolyU(1)_HairpinDistance(10)_Tornado-Corn_ |
| 159498 | pAV-U6+27-Linear-1_PolyU(4)_HairpinDistance(10)nts_Tornado-Corn_ |
| 159499 | pAV-U6+27-Linear-2_PolyU(1)_HairpinDistance(0)nts_Tornado-Corn_ |
| 159500 | pAV-U6+27-Linear-2_PolyU(4)_HairpinDistance(0)nts_Tornado-Corn_ |
| 159501 | pAV-U6+27-Linear-2_PolyU(1)_HairpinDistance(10)nts_Tornado-Corn_ |
| 159502 | pAV-U6+27-Linear-2_PolyU(4)_HairpinDistance(10)nts_Tornado-Corn_ |

| | |
|---|---|
| 159503 | pAV-U6+27-Linear-3_PolyU(1)_Tornado-Corn_ |
| 159504 | pAV-U6+27-Linear-3_PolyU(1)_HairpinDistance(10)nts_Tornado-Corn_ |
| 159505 | pAV-U6+27-Linear-3_PolyU(4)_Tornado-Corn_ |
| 159506 | pAV-U6+27-Linear-3_PolyU(4)_HairpinDistance(10)nts_Tornado-Corn_ |
| 159507 | pAV-U6+27-Linear-3_PolyU(1)_HairpinDistance(0)nts_Tornado-Corn_ |
| 159508 | pAV-U6+27-Linear-3_PolyU(4)_HairpinDistance(0)nts_Tornado-Corn_ |
| 159509 | pAV-U6+27-Linear-3_PolyU(4)_HairpinDistance(14)_Tornado-Corn_ |
| 159510 | pAV-U6+27-Linear-3_PolyU(4)_HairpinDistance(15)nts_Tornado-Corn_ |
| 159511 | pAV-U6+27-Linear-3_PolyU(4)_HairpinDistance(16)nts_Tornado-Corn_ |
| 159512 | pAV-U6+27-Linear-3_PolyU(4)_HairpinDistance(17)nts_Tornado-Corn_ |
| 159513 | pAV-U6+27-Linear-3_PolyU(4)_HairpinDistance(18)nts_Tornado-Corn_ |
| 159514 | pAV-U6+27-Linear-3_PolyU(4)_HairpinDistance(19)nts_Tornado-Corn_ |
| 159515 | pAV-U6+27-Linear-3_PolyU(4)_HairpinDistance(20)nts_Tornado-Corn_ |
| 159516 | pAV-U6+27-Linear-1_PolyU(3)_HairpinDistance(10)nts_Tornado-Corn_ |
| 159517 | pAV-U6+27-Linear-2_PolyU(3)_HairpinDistance(10)nts_Tornado-Corn_ |
| 159518 | pAV-U6+27-Linear-2_PolyU(5)_HairpinDistance(10)nts_Tornado-Corn_ |
| 159519 | pAV-U6+27-Linear-1_PolyU(5)HairpinDistance(10)nts_Tornado-Corn |
